# Eye-tracked Maxwellian illumination system for power-efficient retinal stimulation

**DOI:** 10.64898/2026.09.01.748645

**Authors:** Nathan Jensen, Mohammad Asif Zaman, Linh Vu, Ludwig Galambos, Daniel Palanker

## Abstract

Augmented reality glasses typically have poor optical efficiency (< 1%), which is detrimental for high power applications, such as prosthetic vision. Maxwellian view optics are a candidate for increasing efficiency, but suffer from a very limited exit pupil and thus field of view. Many approaches to extend the eyebox inherently decrease the optical power. We present a design that uses a high-density VCSEL array and eye tracking to create an extended eyebox without sacrificing the advantages of a Maxwellian view display. Our system achieves a 20 degree field of view, a resolution 48.8 LP/mm (10 µm), and an end-to-end optical efficiency of 83%.

## 1. Introduction

The combination of augmented reality glasses and ophthalmic neuroengineering has facilitated a transformative era in vision restoration, enabling innovations such as photovoltaic retinal prostheses [1] and optogenetic therapy [2]. These technologies are based on stimulation of the inner retinal neurons in patients blinded by the loss of photoreceptors due to age-related macular degeneration (AMD) or retinitis pigmentosa (RP).

Photoreceptors are extremely sensitive to light, typically operating at ambient light intensities in the range of nW/mm^2^ on the retina. As such, augmented reality (AR) glasses for normal vision do not generally optimize for optical power. Photovoltaic, optogenetic, photothermal and optoacoustic retinal stimulation, on the other hand, require several orders of magnitude higher light intensity – typically in the range of mW/mm^2^. Such an intense illumination is often limited by ocular safety and inefficient delivery of light to the retina from augmented reality (AR) glasses. In this paper, we present a design of AR glasses to address these concerns. We specifically focus on photovoltaic retinal prosthesis, but similar problems and solutions apply to optogenetic, optoacoustic, photothermal or any other optical methods of retinal activation which require much higher irradiance than ambient light. The techniques can also be used for a high-brightness, high-resolution heads up display.

AR devices typically utilize on Newtonian illumination, which projects a wide beam from an emissive display onto the cornea, providing a large field of view (FOV) (Fig. 1(a)). As a result, most of the optical power is clipped by the pupil, proportional to the FOV of the system (Fig. 1(b)). For example, assuming a uniform beam with a diameter 6 mm on iris entering a 2 mm pupil, only 1/9 of the power is transmitted into the eye. In high power applications, this inherent inefficiency can lead to excessive heating of the AR headset and of the iris. Additionally, in this paradigm, variation of the pupil size with ambient lighting (constriction outdoors vs. dilation indoors) affects the amount of power delivered to the retina. Pupil constriction in bright ambient lighting reduces the amount of power on the retina, dimming the prosthetic vision.

**Fig. 1.**
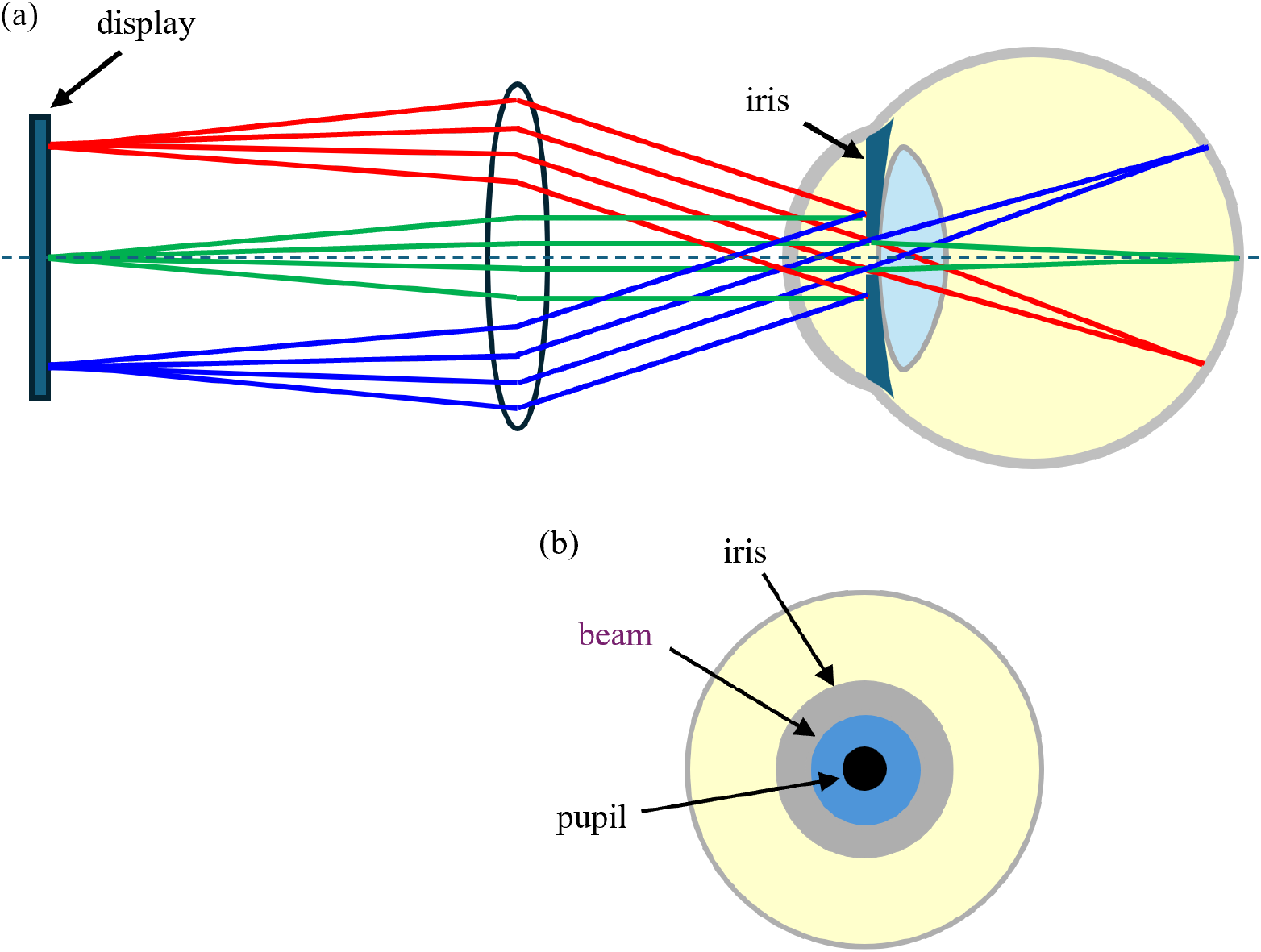
(a) Simplified diagram of the image projection in conventional (wide-field) augmented reality glasses. Light from the pixels of the emissive display is collimated by the lens and imaged onto the retina by the cornea and crystalline lens of the eye. (b) Beam on iris is wider than the pupil, which allows for eye scanning within the range of eye box. However, only a fraction of the beam which passes through the pupil is transmitted onto the retina.

On the other hand, Maxwellian view optics images a light source onto a spot fully contained within the pupil, while the display is imaged onto the retina (Fig. 2). This results in efficient power transmission through the pupil, independent of its size. In addition, the small beam diameter on the eye lens reduces aberrations and provides larger depth of focus. The primary limitation of this technique is its very small exit pupil, or eyebox, which hampers eye scanning as even a small movement of the eye can cause the beam to miss the pupil completely. Moreover, in high power applications, the small beam waist may cause local heating of the iris, which carries safety concerns [3].

**Fig. 2.**
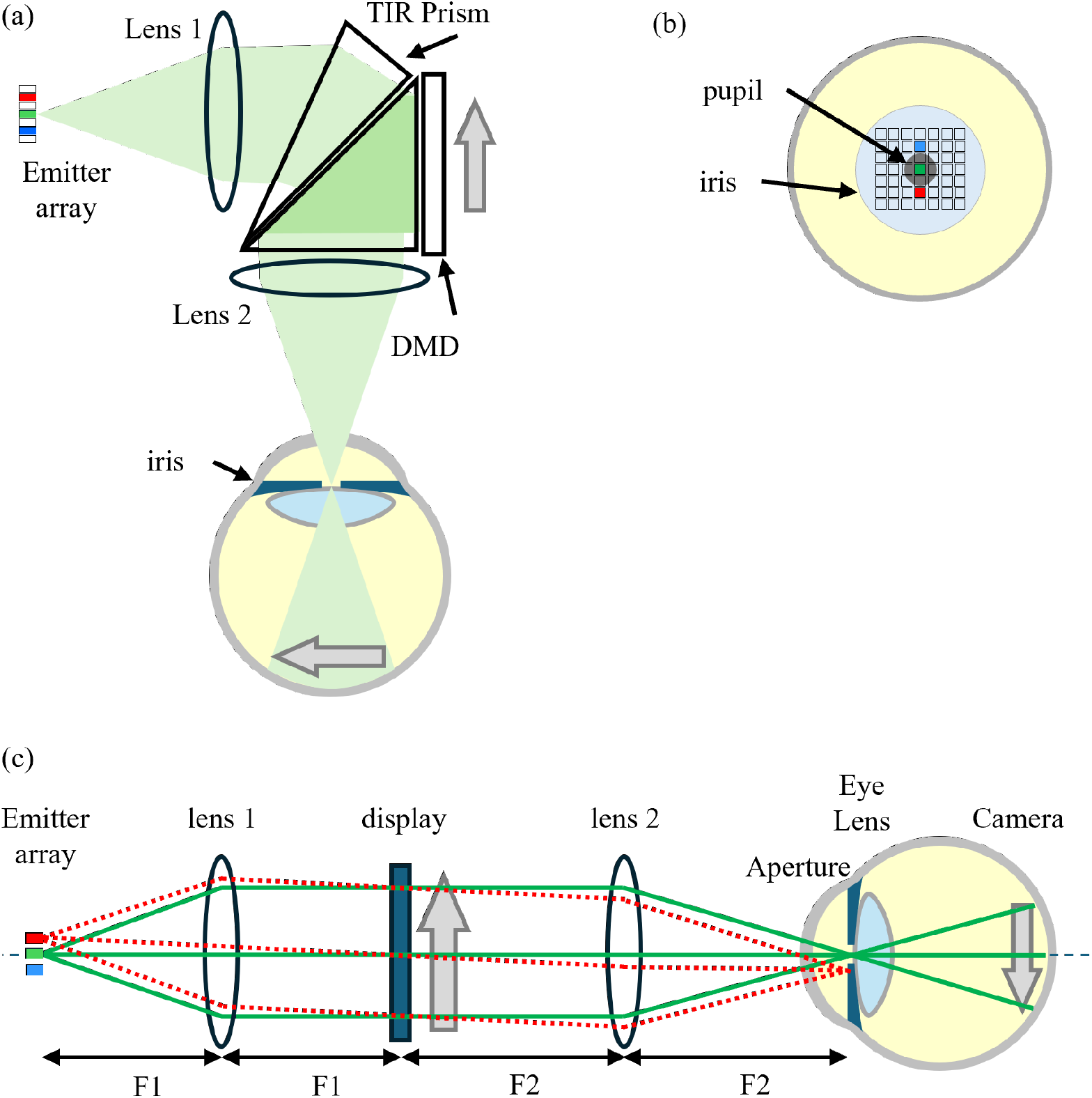
Diagram of a Maxwellian imaging system. a) Light is collimated by lens 1, folded by TIR prism onto the DMD, which forms the stimulation pattern. Reflected light is focused by lens 2, passing through the pupil and forming an image of the stimulation pattern on the retina b) Front view of the iris plane with an image of the emitters array. Central emitter (green) is contained within the pupil, and its beam is transmitted to the retina. Side emitters (red, blue) are blocked by the iris, and should not be activated. c) Unfolded design of the benchtop prototype. Light from each emitter in the array is collimated by lens 1 to illuminate the display, which modulates this illumination. Imaging lens (2) creates an image of the emitters array in the iris plane, while the display is imaged onto the retina. Only the emitters whose beam passes through the pupil are activated, as exemplified here by solid green lines from the central emitter in the array. Beam from the top emitter (red lines, dashed) are blocked by the iris, and therefore should not be activated. Bottom emitter (blue), is shown without corresponding rays, for simplicity.

To overcome the eyebox limitation, Maxwellian displays have employed holographical eyebox duplication, beam splitter arrays, computer generated holography (CGH) with image steering via a spatial light modulator (SLM), and beam-steering with spatially distributed LEDs [4]. Each technique has its own advantages and disadvantages, with a large portion of the current research focused on CGH.

Maxwellian optics can theoretically achieve near-perfect end-to-end light transmission, however most implementations of the expanded eyebox fall far behind the ideal case. For example, a projection system with a conventional LED array designed for optical efficiency [5] fails to capture most of the light due to high divergence of the source emitters, with only 0.03-0.04% end-to-end efficiency; and it has very limited resolution. Amplitude SLM CGH reports less than 1% efficiency [6]. Multiplexed holographical elements [7, 8] and beamsplitters [9] form multiple exit pupils simultaneously, only one of which passes through the pupil at any given time, which greatly reduces the power efficiency – providing a 3×3 grid of exit pupils caps efficiency at 11%. Quarter-waveplate geometric phase prisms improve on this, allowing a theoretical 25% efficiency [10]. Any additional optics required for such systems will further reduce their efficiency.

The efficiency of phase based SLM, such as liquid crystal on silicon (LCoS) displays used for CGH [11], is not reported, but has been proposed as an option specifically for vision restoration applications [12]. However, in such an approach, a significant fraction of power is lost in the higher diffraction orders. Additionally, high computational complexity as well as long response time of LCoS displays remain concerns with this technique, as subretinal prostheses may require frame rates of up to 120 Hz for image multiplexing [13].

While it might be possible to use some of these techniques for retinal stimulation, it is far from trivial and has yet to be demonstrated. Here we present a system with relatively low complexity, which achieves very high optical power efficiency, enables sufficient range of eye scanning, and adheres to the ocular safety limits, while maintaining a small form factor suitable for augmented reality glasses. It utilizes a high resolution VCSEL array as a light source with eye tracking based activation to ensure the entire beam always passes through the pupil, while having intensity low enough large enough to ensure iris safety.

## 2. Methods

The proposed optical system is shown in Fig. 2(a). It consists of a high density array of VCSLEs as the light source, a collimating lens, a total internal reflection (TIR) prism, a digital micromirror device (DMD), and an eyepiece lens. A model eye, which allows capturing the image formed on the “retina” at different directions of gaze, is made of an iris, a lens, and a CMOS camera. To simplify the optics (avoiding custom TIR prisms for DMD illumination) and focus on demonstrating optical efficiency, we replaced the DMD with a transmissive screen, as shown in Fig. 2(c). The only major difference between these designs is the folded light path.

### 2.1 Eye model

In the United States, people having visual field below 20 degrees are considered legally blind, so the image on the retina should exceed this dimension. The image on the retina shifts by distance *s*_*r*_ with eye rotation *φ* according to its focal length *f*_*eye*_: *s*_*r*_ = *f*_*eye* ._ *φ* . *π* /180. At the same time, the pupil shifts by *s* _*p*_ = *d* tan (*φ*), where *d* is the distance between the eye rotation center and the iris plane. Following Gullstrand’s schematic eye [14], the focal length of the relaxed eye is approximately 17 mm and the distance from the corneal apex to the pupil is 3.6 mm. The distance from the corneal apex to the horizontal axis of rotation was measured to be 15.3 mm on average [15]. Therefore, the distance from the pupil to the eye’s rotation center is approximately 12 mm. For eye rotation by *φ* = ±10 degrees (20 degrees field of view), the iris shifts by *s* _*p*_ ≈ ±2 mm, while the image on the retina shifts by *s*_*r*_ ≈ ±3 mm. In other words, 1 degree of the visual field corresponds to 0.2 mm on the iris and 0.3 mm on the retina.

Our model eye includes a 2 mm aperture, representing the most constricted pupil [16], a lens with focal length of 19 mm, and a CMOS camera (Basler model acA2040-90um), which models the retina. These are mounted together on a stage with axis of rotation placed 12 mm behind the aperture to simulate the eye motion (Fig. 3).

**Fig. 3.**
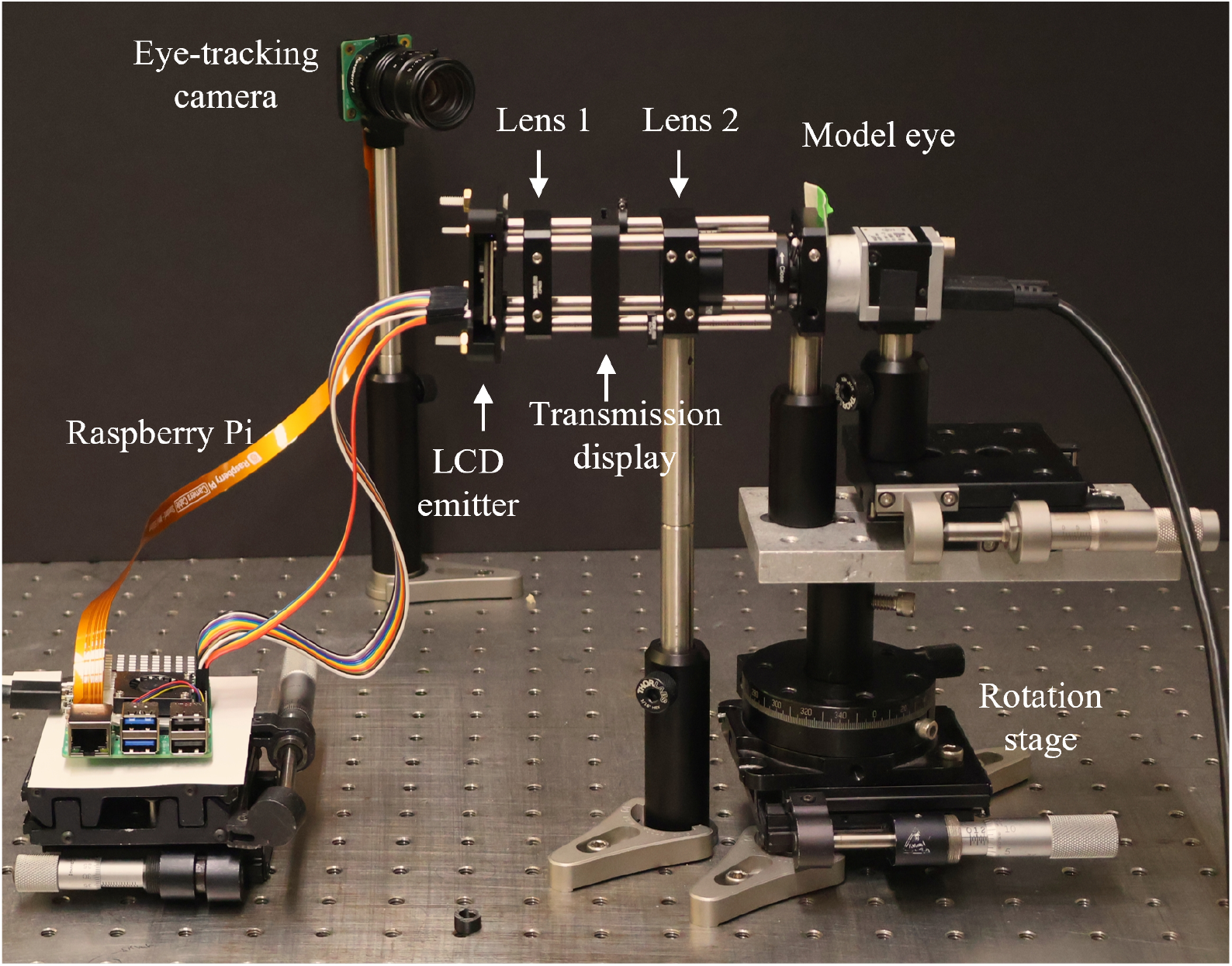
Demo of the optical system. A Raspberry Pi reads eye tracking data and controls the LCD emitter as the light source. Lens 1 collimates the illumination onto a transmission display. Lens 2 images LCD onto the iris of a model eye, which includes a 2 mm pupil and a f=19mm lens imaging the transmission display onto a camera, all mounted on a horizontal rotation stage.

### 2.2 Collimating and eyepiece lenses

For prosthetic vision within 20 degrees of the visual field, the image must be at least 6 mm wide on the retina. For compact AR glasses, the display was sized as, 11 × 8.5 mm matching the dimensions of the DMD (DLP5500, Texas Instruments). Since the image size on the retina, *I*_*ret*_ = *I*_*o*_ (*f*_*eye*_ /*f*_2_), where *f*_2_ is the focal length of the eyepiece lens and *I*_*o*_ is the width of the display, therefore for *I*_*ret*_ = 6 mm, *I*_*o*_ = 11 mm, and *f*_*eye*_ = 19 mm; *f*_2_ ≈ 35 mm.

Collimating lens (lens 1) is a critical component for achieving high optical power efficiency. To maximize power transfer, we matched the collimated beam diameter to the width of the display (*W*_*coll*_ = 11 mm). To collimate a beam with diameter *W*_*coll*_, the lens’ focal distance *f*_1_ = *W*_*coll*_/(2 tan *θ*), where *θ* is the half-angle of the source beam divergence. For *θ* = 17 degrees (typical for a VCSEL array i.e. Osram OLI2020V.A1-850), *f*_1_ ≈ 18 mm.

### 2.3 Light Source

To steer the illumination beam on the pupil, the source must be spatially translatable. As such, the preferred light source is composed of an array of discrete light emitters with a narrow emission cone (≈17 degrees half angle), such as vertical-cavity surface-emitting lasers (VCSELs). Having an array allows the beam to be spatially shifted by turning on and off individual elements rather than their mechanical motion. The pitch, *s*, of the emitter array must be small enough so that for any position of the pupil, a beam of at least one emitter is fully contained within the pupil in its most constricted state (Fig. 4(a,b)). This ensures that the same power is transmitted to the retina at any state of pupil dilation and at any direction of gaze.

**Fig. 4.**
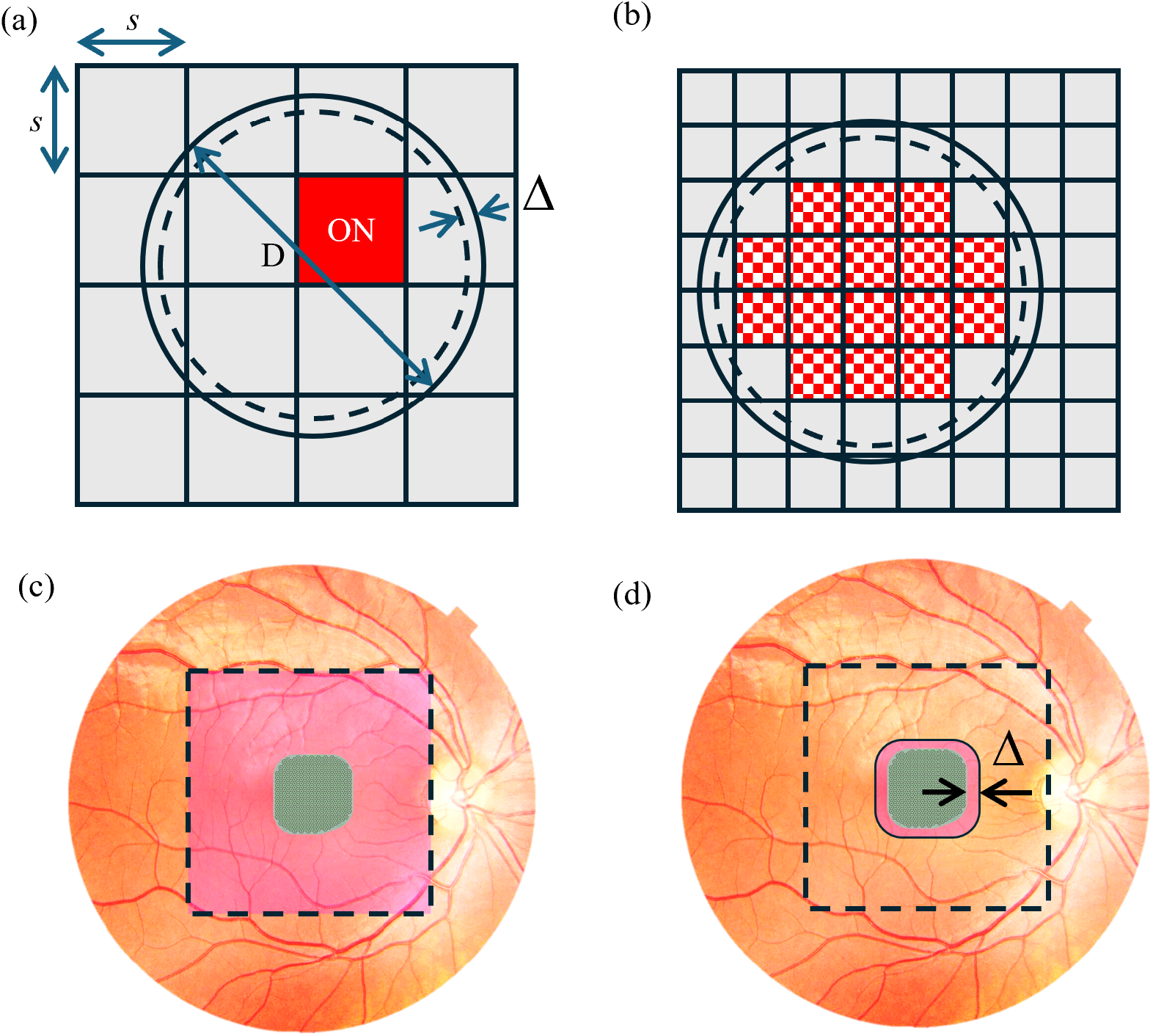
Image of the emitters array in the iris plane. a) A 4×4 emitters array, of a pitch *s*, and a pupil of diameter D. To fit at least one emitter (shaded red, labeled ‘ON’) within the pupil at any position over the array with the eye tracking uncertainty Δ, emitter size should be 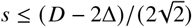. b) With twice smaller emitters covering the same area, multiple emitters fit within the pupil, enabling wider beam (checkerboard shading, red) within the pupil. c) The fundus of a human eye with a 2×2 mm PRIMA implant and 6×6 mm NIR illumination beam. d) By turning off parts of the display that are imaged outside the implant, the illuminated area on the retina is reduced to the size of the implant plus the eye tracking uncertainty margin Δ.

Finally, the total peak power instantaneously available from the source must provide 3.5 mW/mm^2^ irradiance on the retina across the entire image of 6×6 mm, which is 126 mW. To minimize the power density on cornea and iris, the beam on the pupil plane should be matched to the pupil size, or in practice slightly smaller to account for eye tracking uncertainty and resolution. In human eye, the pupil size typically ranges from 3 to 6 mm, depending on a variety of factors including age and lighting conditions, although it has been reported to be as small as 1.8 mm in the most extreme cases [16–18]. We assume a pupil size of >2 mm and require the optical beam diameter on the pupil to be 2 mm for our study. To provide a 2 mm illumination spot over an image shift of ±2 mm (± 10 degrees), the total eyebox size comes to 6 mm. Based on our optical design (about 2x magnification of the emitters on the iris plane), this corresponds to a source array at least 3 mm wide, with a 1 mm section illuminated simultaneously.

A high-density array with individually or matrix addressable emitters having pitch <100 µm allows these conditions to be met in real time when used in conjunction with an eye tracking system. To provide the correct power on the retina, with 100 µm pitch, a 10×10 array of VCSELS is simultaneously activated with a peak power of 1.26 mW per VCSEL, corresponding to a power density of 12.6 W/cm^2^ if the system had a perfect power transmission. This resolution allows pupil tracking in steps of *d*_*t*_ = *s. f*_2_/ *f*_1_ = 100. 35/18 200 ≈ µm, or 1 degree. Several VCSEL arrays reported in the literature meet these specifications [19–22].

Furthermore, with such a high-density display, if the pupil size is also tracked [23], the size of the illumination source can be adjusted to match the pupil diameter. This will ensure the power density on the surface of the eye is kept to a minimum (Fig. 4(b)).

For simplicity of prototyping, in this study we used a TFT LCD display (Adafruit Mini PiTFT 1.3”) with a pitch of 108 µm instead of a customized VCSEL array to demonstrate to demonstrate the effect of eye rotation with and without pupil tracking based real-time beam steering. It should be noted that all optics after the collimating lens function the same, so only the illumination source differs. However, due to the wide emission of the LCD array emitters, the power efficiency would be significantly lower than with VCSELs. Therefore, for power efficiency measurements, we used a VCSEL emitter (Osram V100P000A-680).

To ensure that the image has a constant brightness, the illumination beam on the display must be uniform and speckling must be kept to a minimum. VCSELS typically have Laguerre-Gaussian modes in their radiation profile, which changes based on the drive current [24]. A VCSEL array that has size and specs similar to those required for this system achieves a reasonably flat beam profile, with the center maintaining 75% of the peak optical power (Osram OLI2020V.A1-850A). One way to increase the beam uniformity is to mix different parts of the beam on a display using a homogenizer i.e. fly’s eye [25]. Beam shaping with a diffractive optical element or meta optics are other options that preserve optical efficiency – for single mode VCSELS with a Gaussian emission, a top-hat beam shaper could be used.

The minimum speckling contrast, *C*_*min*_ depends on the number of emitters, *n*, as 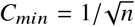 [26]. With 100 VCSELs the minimum speckling contrast drops 10 fold compared to a single VCSEL.

### 2.4 Iris Safety

Illumination for retinal implants, at 3.5 mW/mm^2^ and 30% duty cycle [27], adds up to average irradiance of 1.05 mW/mm^2^, which is well below the safety limit of 4.6 mW/mm^2^ for 880 nm radiation [3]. However, when the entire beam of 6 × 6 mm on the retina is concentrated into a small spot at the pupil plane, irradiance may exceed the iris safety limit. Since the illumination of retinal areas outside the implant can be turned off, as illustrated in Fig. 4, the total power in the beam waist on the pupil will be reduced accordingly. To guarantee a complete coverage of the implant, the illuminated zone should exceed the implant size by uncertainty of eye tracking. For example, Δ = ±1 degree of precision in eye tracking corresponds to ±0.3 mm on the retina.

Fig. 4(d) demonstrates an illuminated area of 2.6 × 2.6 mm centered on a 2 × 2 mm implant with ±0.3 mm margins beyond the implant. In this case, the total illuminated area on the retina decreases from 36mm^2^ in Fig. 4(c) to just 6.8 mm^2^. This area corresponds to a total peak power *P*_*r*_ = 3.5 mW/mm^2^ × 6.8 mm^2^ ≈ 24 mW, and with a 30% duty cycle the average power is < *P*_*r*_ >= 24 ∗ 0.3 ≈ 7.2 mW. In the 2 mm diameter beam on the pupil illustrated in Fig. 4(b), the average irradiance is about 2.3 mW/mm^2^ – which is below the safety limit.

For optogenetic stimulation, similar considerations apply, but the illuminated area on the retina should correspond to the transfected zone, typically limited to the perifoveal ring [28].

### 2.5 Eye Tracking

Eye tracking was performed using a Rasberry Pi 5 microcomputer with the Rasberry Pi Global Shutter Camera. Tracking was implemented in a python script using a center of mass algorithm [29] to track a target mounted on the model eye with a precision of < 1 degree. The Rasberry Pi then controlled the illumination array (i.e., LCD display) based on the tracking data, allowing real time eyebox positioning.

### 2.6 Transmission Display

A transmission display was created using 200nm thick titanium patterns lithographically fabricated on quartz wafers by e-beam evaporation process. Lift-off was enabled by a bilayer photoresist: a non-photosensitive underlayer (Shipley LOL-2000) followed by a photosensitive imaging layer (Shipley SPR-3612). Patterns were exposed with the 405nm laser of a Heidelberg MLA150 direct write lithography tool. After the metal deposition, the bilayer photoresist was removed by soaking the wafer in Shipley 1165 Remover overnight, heating up at 80° C for 1 hour, with a subsequent sonication and a solvent clean.

## 3. Results

We measured the key optical parameters of the assembled benchtop prototype (Fig. 3), including the exit pupil size, image size and quality on the retina, instantaneous and extended eyebox size, and finally the end-to-end power efficiency of the system.

The exit pupil size was measured by replacing the eye model with a diffusive screen placed at the pupil plane. The beam on the screen was then imaged with a camera, using a ruler placed nearby as a scale. The beam waist, representing an image of the LCD array in the pupil plane, was measured (Fig. 5(a)) using the captured image. Based on the 108 µm pixel pitch of the source display, a 9 × 9 pixel square should correspond to 1.9 × 1.9 mm in the pupil plane. We measured the square to be 1.8 × 1.8 mm, close to our estimations. In practice, a circular source would be used to better fill the pupil, which is also shown in Fig. 5(a).

**Fig. 5.**
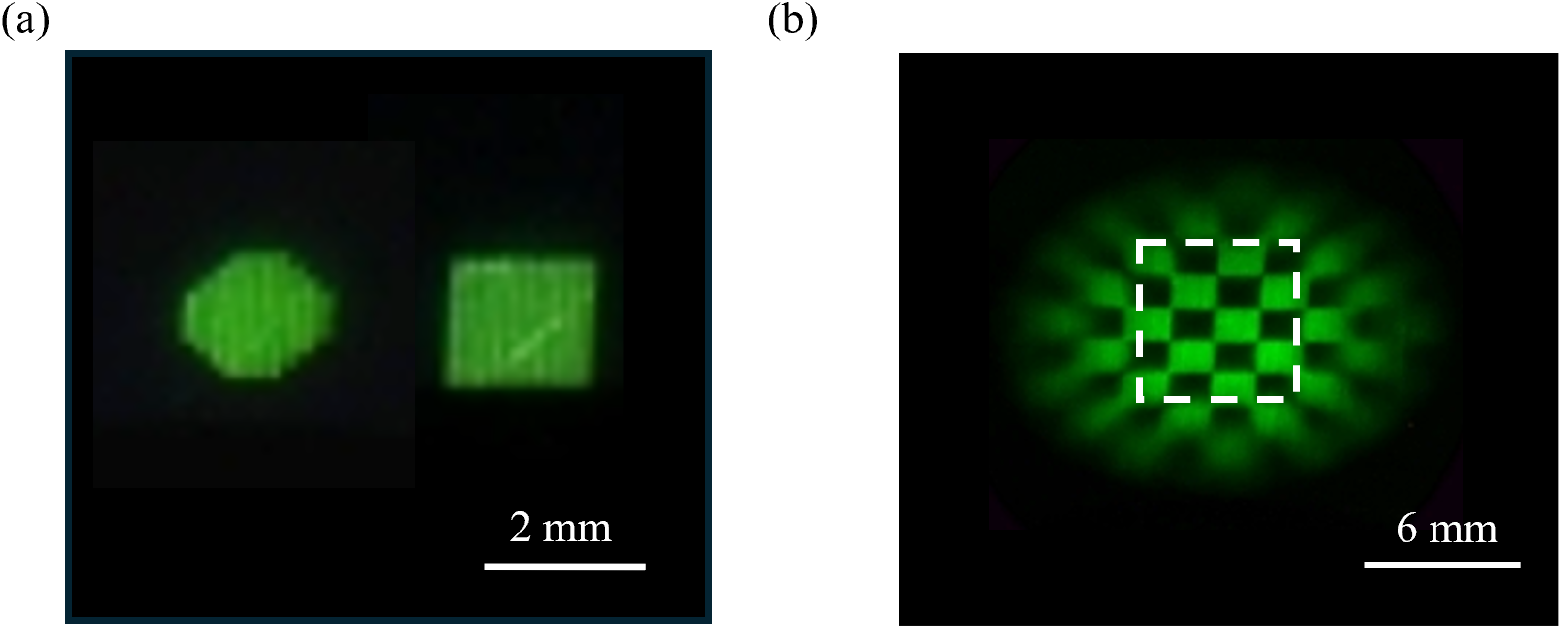
a) Images of the 9 × 9 pixel square and 9-pixel diameter circle projected by the LCD emitter array onto the iris plane, where it is roughly a 1.8 × 1.8 mm in size. b) Checkerboard pattern, composed of 9 × 9 pixel squares shows a 14 mm width of the extended eyebox, though severe attenuation and aberration affect the edges. A 6 mm square region marked by white dash line represents the 20 degrees eye box.

The common practice for characterizing the eyebox of similar systems is to measure the width of the emitter array on the pupil plane. As shown in Fig. 5(b), our eyebox measured 14 mm in width. In principle, this linear dimension can be increased using a larger source array and lenses. However, in reality, such extension does not necessarily correspond to widening the visual field for an eye rotating by large angles. Therefore, we also measured the “rotational eyebox” – changes in brightness of the retinal image with eye rotation – with and without the eye-tracking-based shift of the light source. Without eye tracking, rotation by more than 2° caused dimming by > 10%. With eye tracking, dimming was < 10% throughout ±10 degrees of eye rotation. In fact, the image remained visible even with rotations as extreme as ±23 degrees, although aberrations and severe narrowing of the apparent aperture became very significant. Parts of the image exhibited 50% dimming at ±15 degrees, corresponding to a rotational eyebox size of 8 mm. This shows that due to mismatch between the source translation and eye rotation planes, the aberration and attenuation-free eyebox is significantly smaller than predicted by a linear translation alone.

The image size on the retina was calibrated using the 5.5 µm pixel pitch of the model eye camera. The transmission display representing the DMD was 11 × 8.7 mm and its image of 1050×821 pixels represents 5.8 × 4.5 mm, corresponding to a magnification of 0.52 × . The design magnification was 0.54 ×, with the corresponding image of 6 × 4.75 mm. The very small deviation is likely due to lenses having slightly different focal lengths and imperfect alignment.

The image quality was analyzed using USAF 1951 object to find the maximum resolvable spatial frequency using the Rayleigh criterion. USAF 1951 group four element five was resolvable with a contrast of 33% (Fig. 6). Based on the image magnification of 0.52, on the camera representing a human retina, this corresponds to 48.8 LP/mm, 14.7 LP/degree, or approximately 10 µm resolution – about half the level of normal vision.

**Fig. 6.**
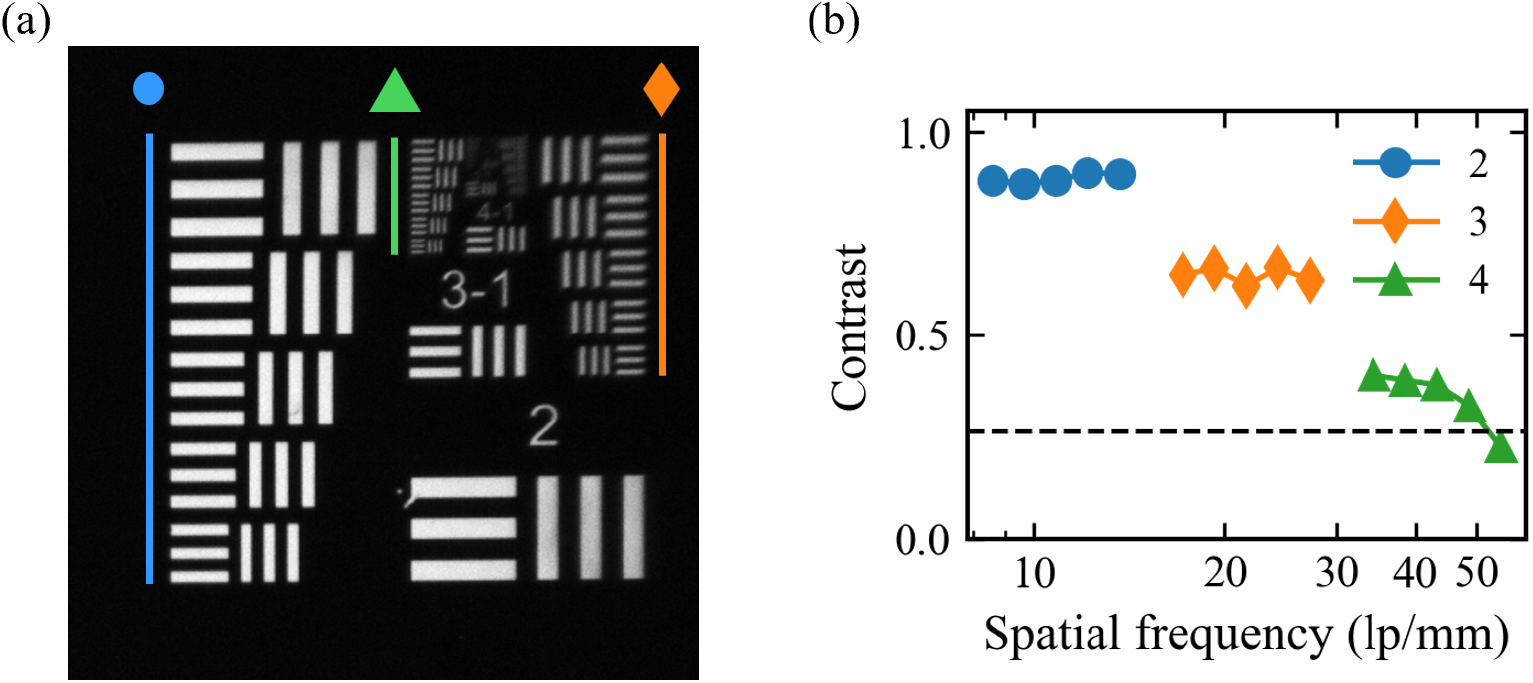
a) Resolution test using USAF 1951 target image as seen by the eye model’s camera, zoomed in to groups 2-8. Lines and symbols mark group 2 (circle), 3 (diamond), 4 (triangle). b) The contrast of groups 2-4, elements 2-6, with the Rayleigh threshold of 26.4% shown as a horizontal dash line. The final point above the threshold corresponds to a spatial frequency of 48.8 lp/mm.

We further evaluated the optical fidelity of our system with a variety of images shown in Fig. 7: including symbols (Fig. 7(a,c)) and text filling a 20 degree visual field (Fig. 7(b)). Fig. 7(d-f) illustrate the technique of limiting the image size on the retina to that of the implant to reduce the power density in the pupil plane (diagrammatically shown in Fig. 4(d)). For this demo, we used windowed blocks of text. The eye was rotated by 10 degrees left (Fig. 7(d)) and right (Fig. 7(f)), such that it is centered on different portions of the text, as done naturally while reading. Areas outside of the 2 × 2 mm central region, which would fall outside the retinal implant, are blacked out. Eye-tracking-based shift of the light source on LCD maintained the retinal illumination, although reflections and aberrations by the lens of the model eye cause some attenuation and reduced resolution at the edges of the visual field when the eye was rotated.

**Fig. 7.**
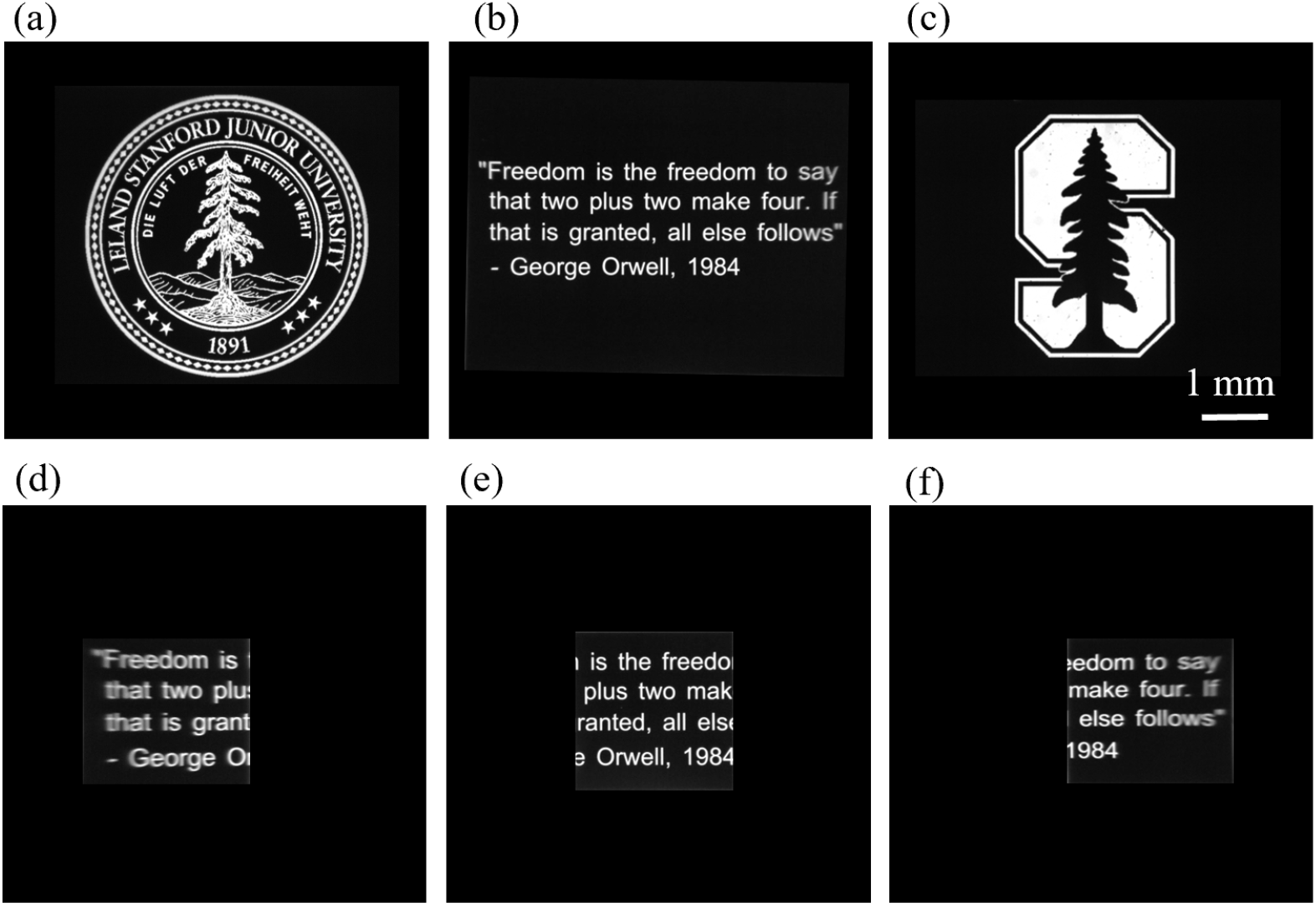
a) University Seal demonstrating high resolution of the imaging system. b) Text filling roughly 20 degrees of the visual field, or 5.8 mm on the camera. c) University logo used for showing the eyebox expansion with active eye tracking in Fig. 8 and Supp. Video 1. The 2 × 2 mm portions of the text are shown, while the rest of the display is “turned off” to minimize the power density on the pupil, as illustrated in Fig. 4(d). The camera rotation is at +10 degrees (d), 0 degrees (e), and −10 degrees (f).

Eye tracking-based correction of the light source is further illustrated in Fig. 8. With a shifting light source, images in the eye rotated by 5 and 10 degrees have the same brightness (Fig. 8(c)). This is in stark contrast to not using eye tracking, where the image is severly attenuated at 5 degrees and nearly imperceptible at 10 degrees of rotation (Fig. 8(b)). The white arrow in Fig. 8(a) points at the 1 mm green spot on LCD, which moves to compensate for eye rotation. Videos demonstrating this process across the entire ±10 degree rotation are provided as Supplementary Video 1 and 2.

**Fig. 8.**
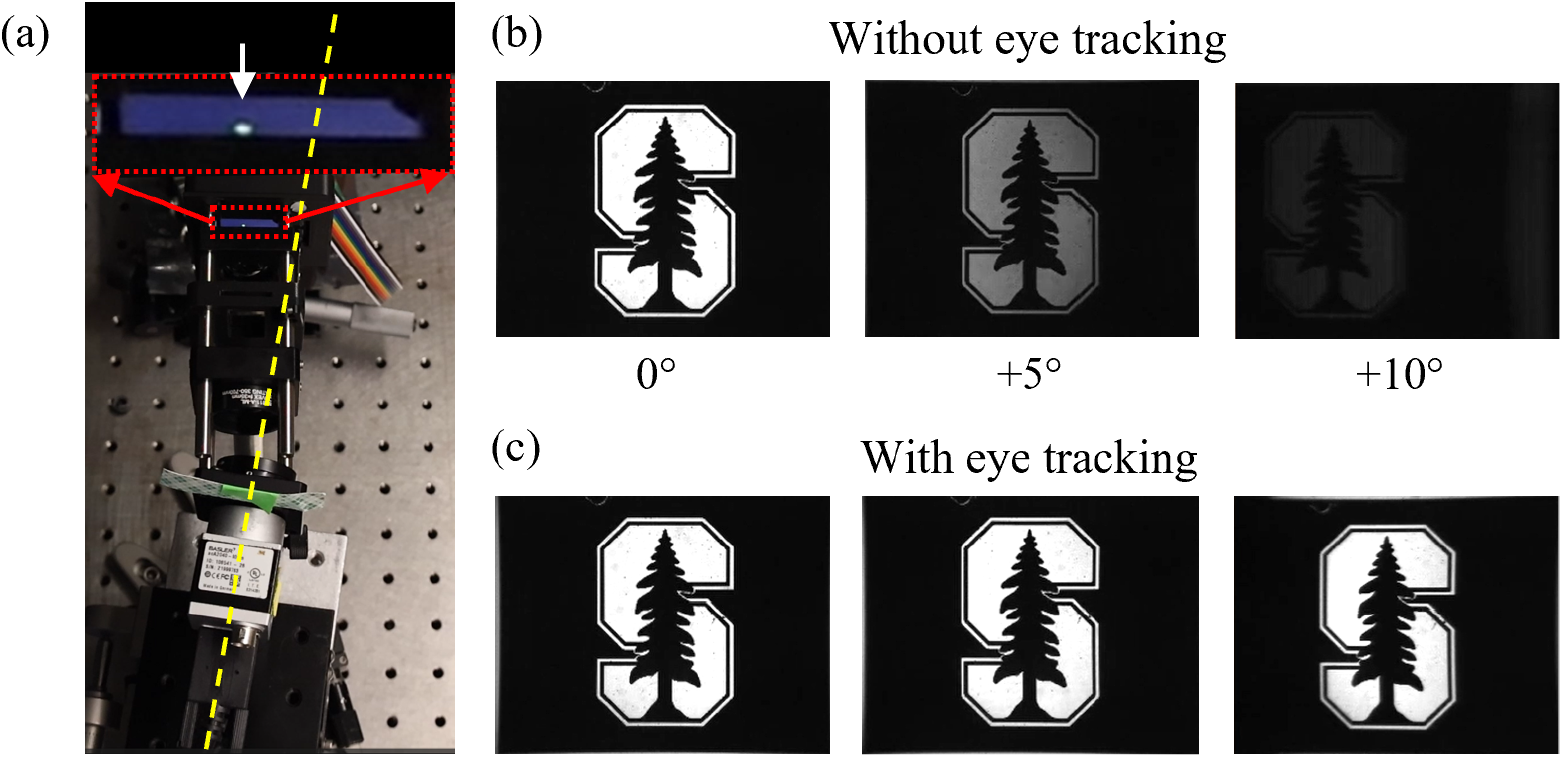
Demonstration of the eyebox expansion via real time eye-tracking. a) Top-down view of the setup with the eye model rotated by +10 degrees, indicated by yellow dashed line. At the top is a zoomed inset (red dotted lines) of the source LED display, with an arrow pointing at the light emitting area. b). Camera captured images of the object shown in Fig. 7(c) when the eye model is rotated by 0, 5, and 10 degrees, and no tracking or eyebox compensation is used. c) The same view as in (b), but with eye tracking and real-time beam steering for eyebox expansion. The real-time videos showing a full ±10 degrees rotation with and without eye tracking and eyebox expansion are provided in Supp. Video 1 and 2.

Finally, the end-to-end power efficiency was measured using a power meter (OPHIR 1Z01600 and PD300-TP) by comparing the power at the source to the power measured beyond the pupil. We recorded 10.3 mW optical power from the VCSEL source, 9.0 mW after lens 2, and 8.53 mW before the camera sensor of the eye model. This yields an end-to-end power efficiency of 83%.

## 4. Discussion

We presented a design of a power-efficient augmented reality system for high-power retinal stimulation, based on Maxwellian view optics and eye tracking for expanded eyebox. Performance of our benchtop prototype came very close to the design targets, with optical power transmission efficiency of 83% using off-the-shelf optical components. A majority of this power loss is accounted for by reflections from the 3 lenses. While the design implemented here used a transmissive screen as the object for image projection, it is very likely that augmented reality glasses would use a DMD, LCoS, or a similar reflective display. These components typically have efficiency peaking around 70% [30], which would bring down the overall efficiency to 58%. To the authors’ best knowledge, this is still much higher than the end-to-end efficiency of Maxwellian systems featuring eyebox-expansion reported in the literature to date, all of which are below 1%, except for a claim of 25% efficiency for eyebox expansion, ignoring all other optics [10].

These estimates do not include a potential power loss due to overfilling the rectangular display if a round beam exceeds its dimensions; which can be avoided by underfilling the display. Another aspect not included in this study is a beam homogenizer between the light source and a display since its design depends on actual implementation of the VCSELs array, not available off-the-shelf. These effects are also not taken into account by most power efficiency values reported in the literature.

The real time eye tracking for eyebox expansion eliminates one of the major downsides of a Maxwellian view display. In the case of retinal prostheses, this allows natural “eye scanning” over a 20-degree field of view, which should significantly improve convenience of the PRIMA system, currently limited to eye scanning within a pupil diameter [1, 31]. In principle, this system can provide a wider field of view for other augmented reality applications.

Furthermore, eye tracking combined with pupil size monitoring allows minimizing the power density on the surface of the eye by 1) changing the size of the light source so that the beam size matches the pupil diameter and 2) turning off portions of the display that are not projected onto the implant. These adjustments are important for operating within ocular safety limits.

Finally, this system combines simple optics and software algorithms, in stark contrast to other systems which may accomplish such high efficiency, but require complex optics and even a GPU to enable real time operation, which has not yet been demonstrated [11].

## 5. Conclusion

The use of AR glasses for prosthetic vision faces unique challenges compared to standard applications of AR. Much higher power requirements than for natural vision, necessitates high power efficiency to prevent overheating of the glasses while minimizing their size, as well as careful consideration of tissue safety. The Maxwellian view display with eye tracking presented here is uniquely suited for this purpose. We have demonstrated its ability to provide high power efficiency and good image quality within 20 degrees of visual field, which exceeds the limits of legal blindness.

## Data Availability

Data underlying the results presented in this paper are not publicly available at this time but may be obtained from the authors upon reasonable request.

